## Supplementary material for "Antagonistic behavior of brain networks mediated by low-frequency oscillations: electrophysiological dynamics during internal–external attention switching"

| Patient No. | Age (years) | Gender | Handedness | Suspected seizure onset zone | Epilepsy duration (years) |
|---|---|---|---|---|---|
| P1 | 31 | M | R | Left frontal lobe–premotor area | 2 |
| P2 | 26 | F | R | Right frontal lobe, insula | 13 |
| P3 | 38 | F | R | Right insula, operculum | 9 |
| P4 | 41 | F | R | Right frontal lobe–prefrontal area | 32 |
| P5 | 15 | M | R | Left frontal and temporal lobe | 7 |
| P6 | 16 | M | L | Left insula | 11 |
| P7 | 44 | F | R | Right parietal lobe | 31 |
| P8 | 48 | F | R | Right temporal lobe | 31 |
| P9 | 12 | F | R | Left frontal lobe–prefrontal area | 9 |
| P10 | 15 | F | R | Left temporal lobe | 2 |
| P11 | 43 | M | R | Right temporal, parietal, and occipital lobes | 34 |
| P12 | 46 | M | L | Left frontal lobe | 34 |
| P13 | 29 | F | R | Right frontal and temporal lobe | 20 |
| P14 | 35 | F | L | Left temporal lobe | 21 |
| P15 | 55 | M | R | Right frontal lobe | 27 |
| P16 | 46 | F | R | Right frontal lobe–medial cortex | 44 |
| P17 | 37 | F | L | Right temporal lobe | 13 |
| P18 | 44 | M | R | Left frontal lobe–premotor area | 36 |
| P19 | 36 | M | R | Right temporal lobe | 12 |
| P20 | 38 | F | R | Right frontal lobe | 20 |
| P21 | 32 | M | L | Right frontal lobe–orbitofrontal area | 23 |
| P22 | 28 | F | R | Right frontal lobe | 10 |
| P23 | 24 | F | L | Right temporal lobe | 6 |
| P24 | 49 | M | R | Right temporal lobe | 27 |
| P25 | 30 | F | L | Left temporal lobe | 19 |

**Supplementary Table 1. Subject information.** F–female, M–male, R–right, L–left.

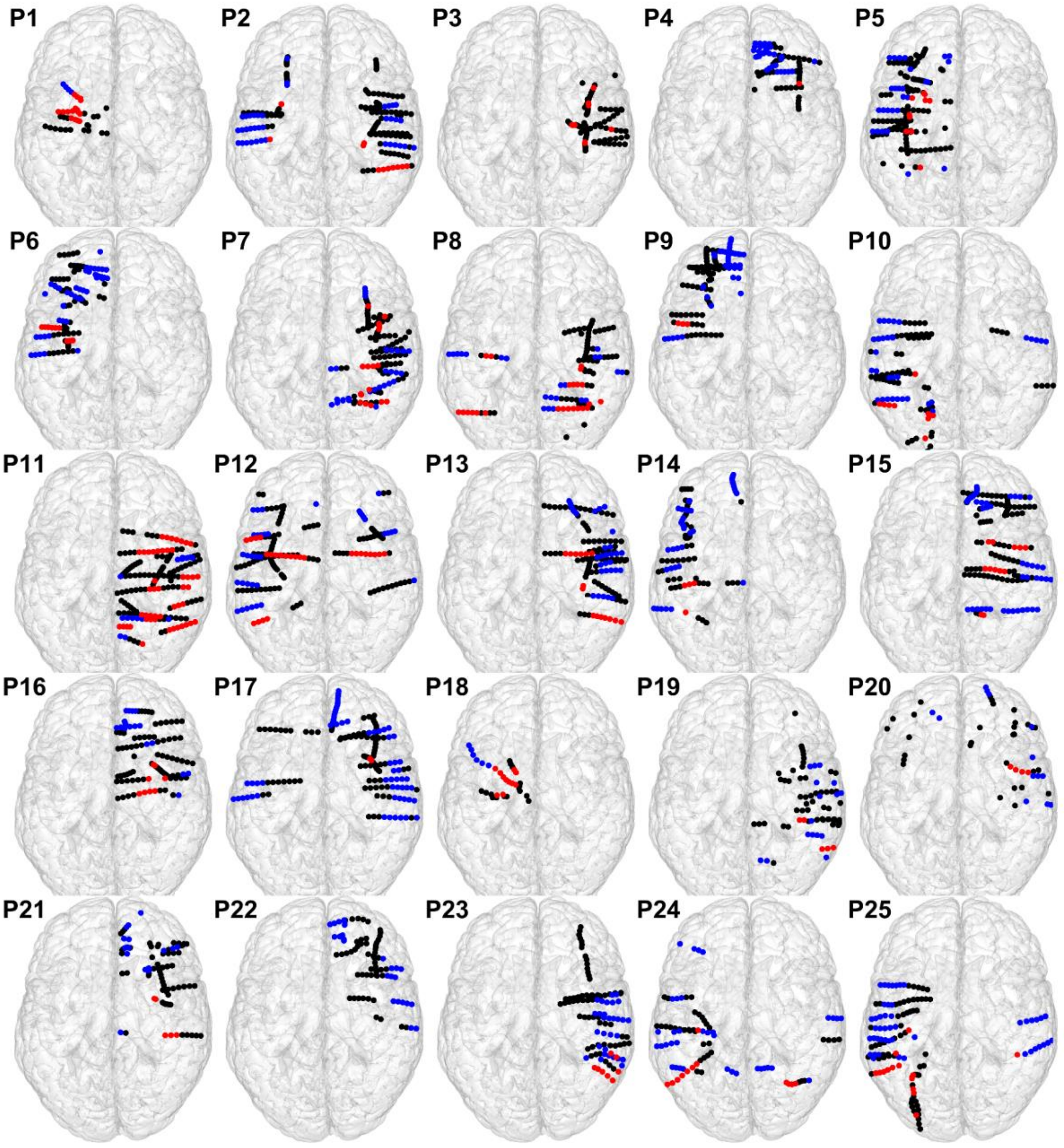


**Supplementary Figure S1. Single-subject (P1–P25) iEEG implantation schemes.** Top views of the MNI brain template (colin27). The colored circles represent the iEEG channels (i.e., the centers of the bipolar-referenced SEEG contact pairs). Note that (1) the iEEG channels were projected up front for visualization purposes (in reality they were located deep inside the brain); (2) some electrodes may appear curved due to MNI normalization (in reality, they were straight); and (3) the size of the circles does not match the physical size of the contacts, which were much smaller (cylindrical shape: 0.8-mm diameter and 2-mm height). The iEEG channels (black circles) were assigned to the DMN (blue) or the DAN (red). Although the implantations were very heterogeneous among subjects, 24 of them had at least some channels in both networks.

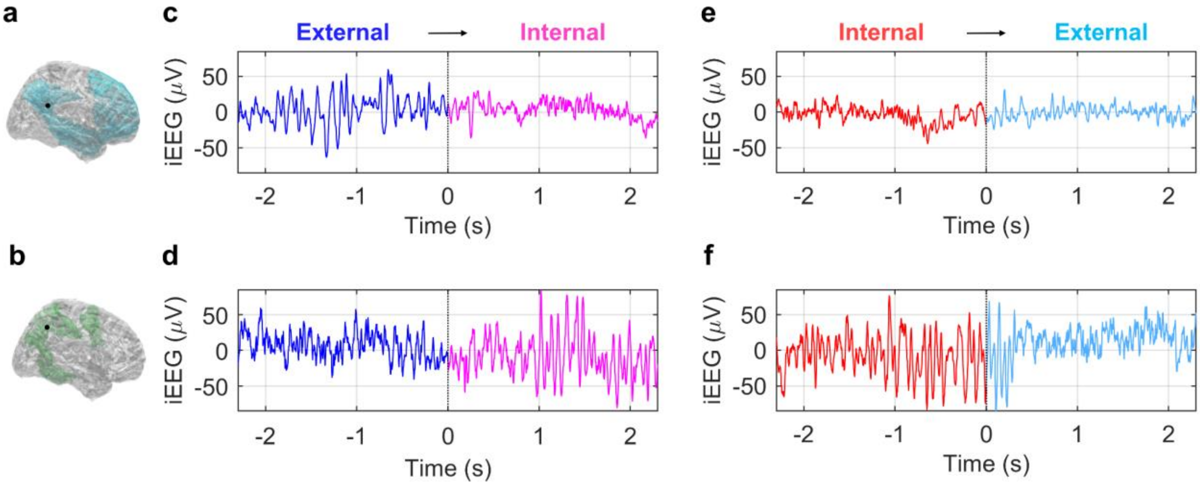


**Supplementary Figure S2. Exemplary single trials of raw iEEG in the DMN and DAN.** Two trials (E-I and I-E) measured simultaneously by two selected channels (from the DMN and DAN, the same channels as in Fig. 2) from subject P8 were selected for an illustration. **(a)** Right lateral right view of the brain template with a highlighted channel position (black dot) and DMN (cyan). **(b)** Same as in A, but for a channel in the DAN (in green). **(c)** A raw iEEG trace of a single trial recorded from the DMN channel during external–internal attention switching. **(d)** The same trial but recorded from the channel located in the DAN. **(e, f)** Another single trial during internal-external attention switching. The same notations as in c and d. The DMN was deactivated during the E-task (the iEEG oscillatory activity was dominated by slower frequency rhythms) and activated during the I-task (the iEEG activity was dominated by faster frequency rhythms). At the same time, the opposite situation was observed in the DAN (activated during the E-task and deactivated during the I-task).

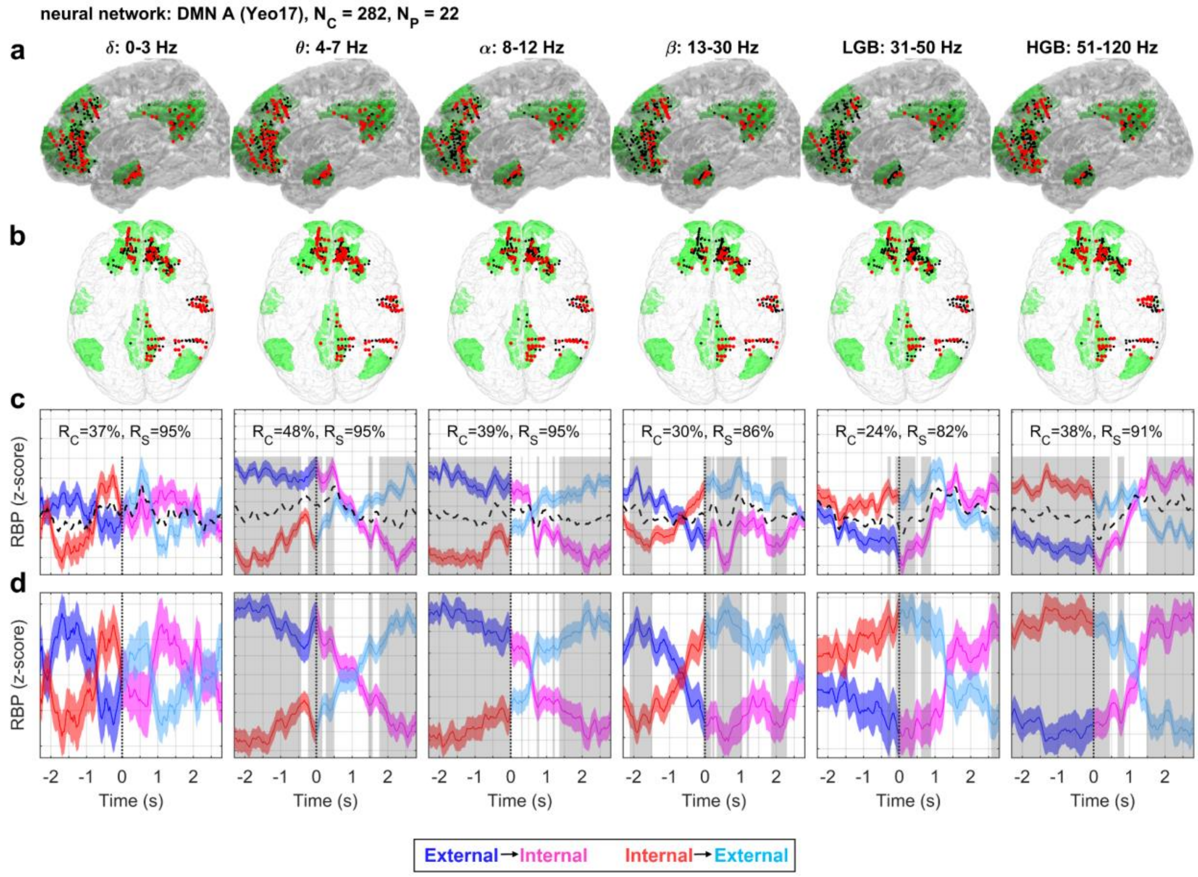


**Supplementary Figure S3. The DMN-A during the attention-switching task.** Same notations as in Fig. 4. The DMN-A (based on the Yeo-17 brain parcellation, highlighted in green) consisted of 282 channels in 22 subjects. The channels were anatomically distributed mostly in the frontal lobe (anterior cingulate cortex, middle frontopolar gyrus, superior and middle frontal gyrus, and rostral gyrus) but also in the temporal lobe (middle temporal gyrus) and parietal lobe (angular gyrus and precuneus). Consistent with overall DMN behavior, there was a pronounced difference in the LFBs (< 30 Hz), especially in the alpha band (8–12 Hz), and we observed the opposite activation in the HGB. The relative decrease of the LFB power and increase in the HGB during the I-task and subsequent switch of the activation during the E-task, suggest DMN-A activation for the I-task (red and magenta curves) and deactivations/suppressions during the E-task (blue and cyan curves).

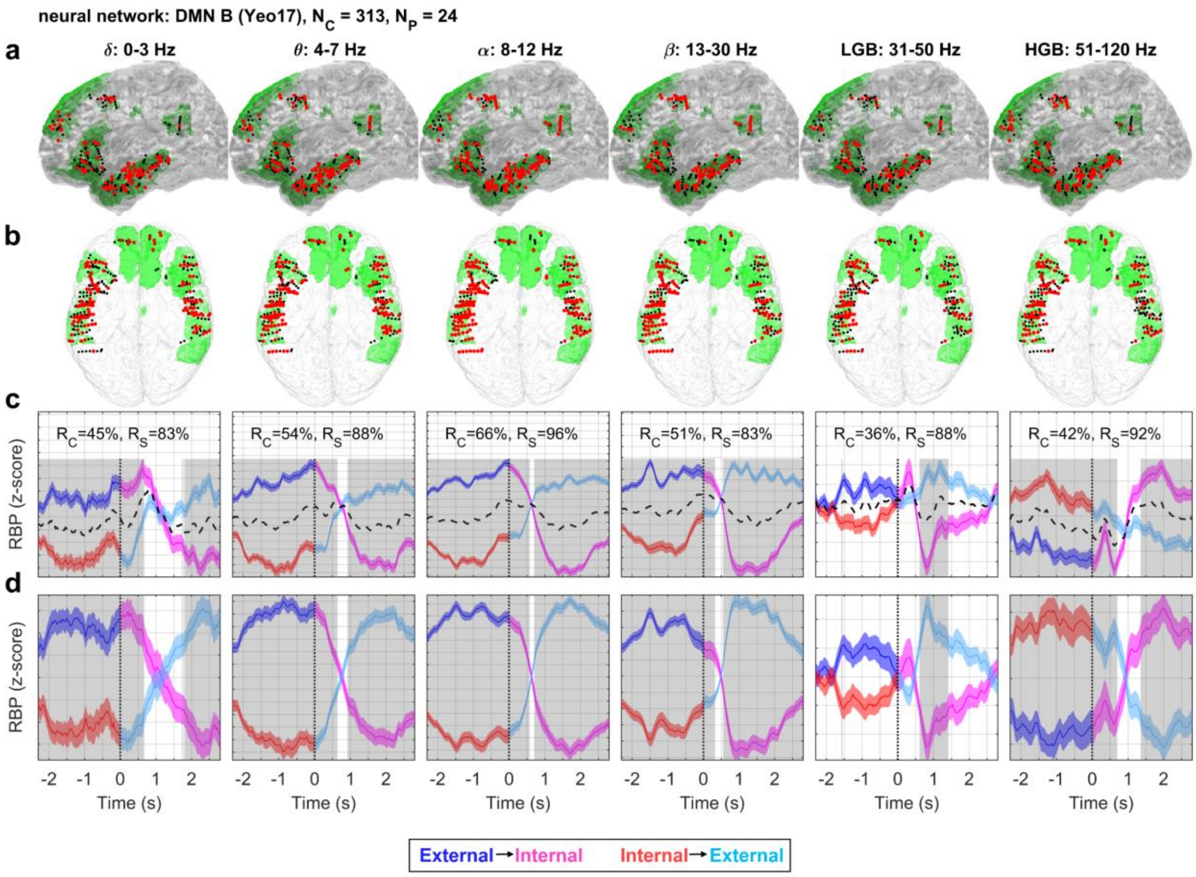


**Supplementary Figure S4. The DMN-B during the attention-switching task.** Same notations as in Fig. 4. The DMN-B (based on the Yeo-17 brain parcellation, highlighted in green) consisted of 313 channels in 24 subjects. The channels were anatomically distributed mostly in the temporal lobe (middle and superior temporal gyrus) but also in the frontal lobe (superior and inferior frontal gyrus) and parietal lobe (angular gyrus). Consistent with overall DMN behavior, there was a pronounced difference in the LFBs (< 30 Hz), especially in the alpha band (8–12 Hz), and we observed the opposite activation in the HGB. The relative decrease of the the LFB power and increase in the HGB during the I-task and subsequent switch of the activation during the E-task, suggest the DMN-B activation for the I-task (red and magenta curves) and deactivations/suppressions during the E-task (blue and cyan curves).

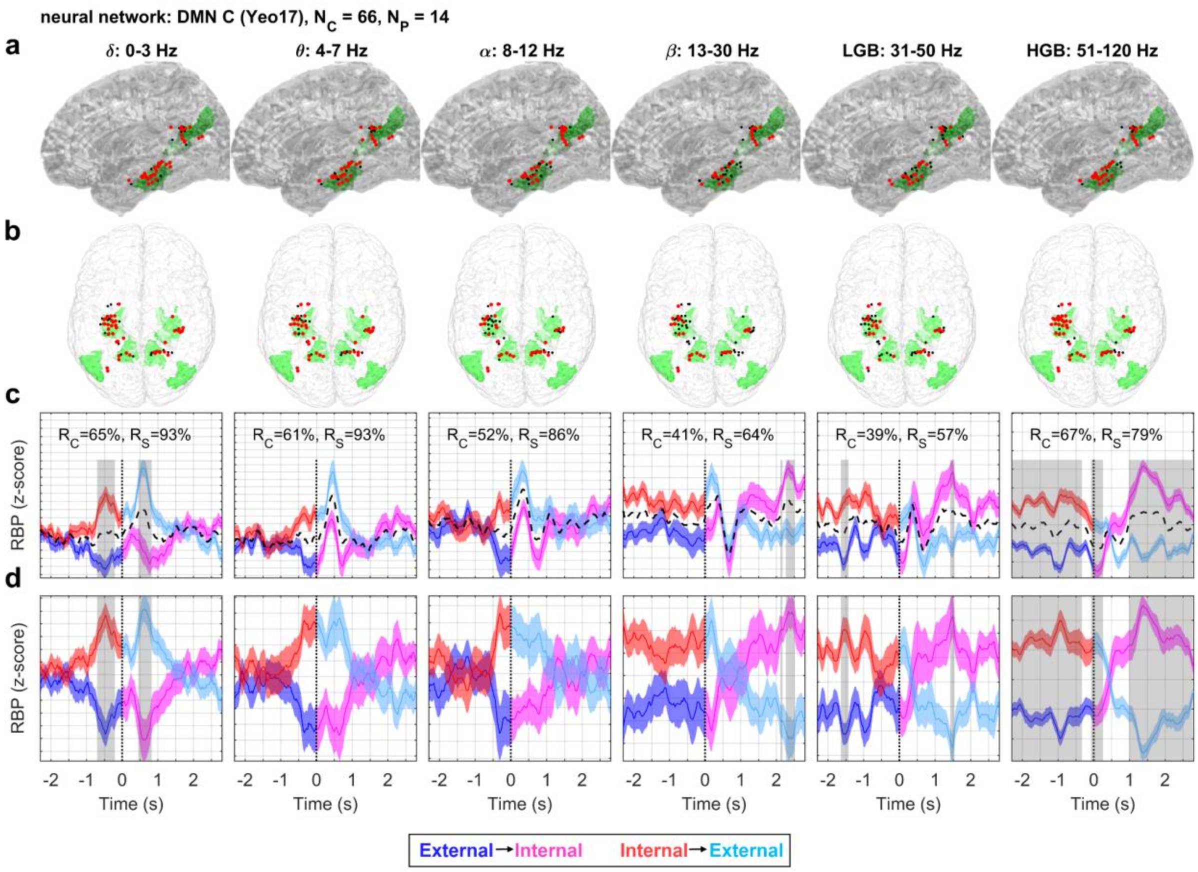


**Supplementary Figure S5. The DMN-C during the attention-switching task.** Same notations as in Fig. 4. The DMN-C (based on the Yeo-17 brain parcellation, highlighted in green) consisted of 66 channels in 14 subjects. The channels were anatomically distributed mostly in the medial temporal lobe (parahippocampal gyrus, fusiform gyrus, collateral sulcus, and hippocampus) but also in the parietal lobe (angular gyrus and posterior cingulate cortex). The RBP was less smooth, presumably due to the lower number of channels. The activity was consistent with overall DMN behavior only in the HGB. In the LFBs, there was a different activity pattern, most evident in the theta band: a decrease of the LFB power before the presentation of the new stimulus ($t$ = -0.5 s) for the E-task (blue curve) and an increase for the I-task (red curve) shortly before the completion of the self-episodic memory retrieval task. However, these differences were not considered significant on the network level ($P$ > 0.001, FDR corrected). The increase in HGB during the I-task and subsequent switch of the activation during the E-task, suggest the DMN-C activation for the I-task (red and magenta curves) and deactivations/suppressions during the E-task (blue and cyan curves).

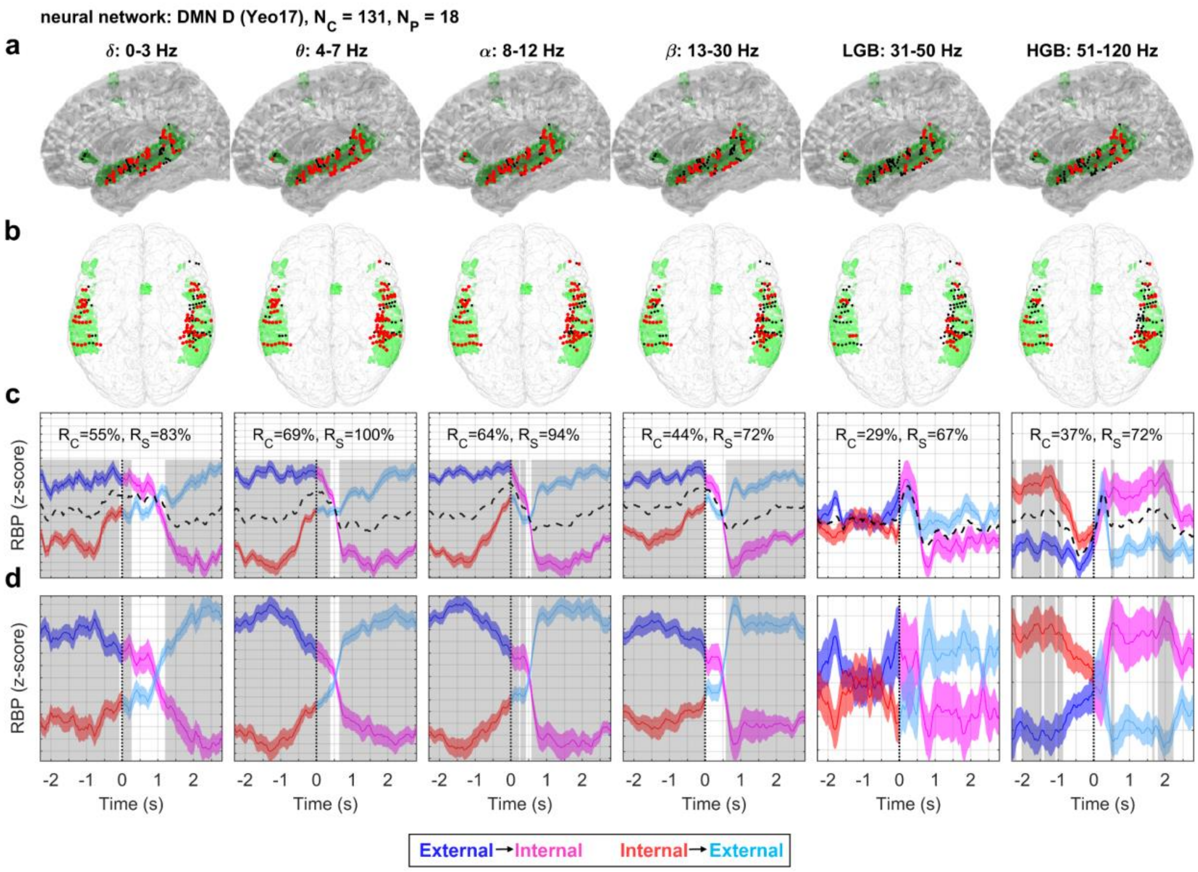


**Supplementary Figure S6. The DMN-D during the attention-switching task.** Same notations as in Fig. 4. The DMN-D (based on the Yeo-17 brain parcellation, highlighted in green) consisted of 131 channels in 18 subjects. The channels were anatomically distributed dominantly in the temporal lobe (superior and middle temporal gyrus), with a few channels in the frontal lobe (inferior frontal gyrus). Consistent with overall DMN behavior, there was a pronounced difference in the LFBs (< 30 Hz), especially in the alpha band (8–12 Hz) and the opposite activation in the HGB. The relative decrease of the LFB power and increase in the HGB during the I-task and subsequent switch of the activation during the E-task, suggest the DMN-D activation for the I-task (red and magenta curves) and deactivations/suppressions during the E-task (blue and cyan curves).

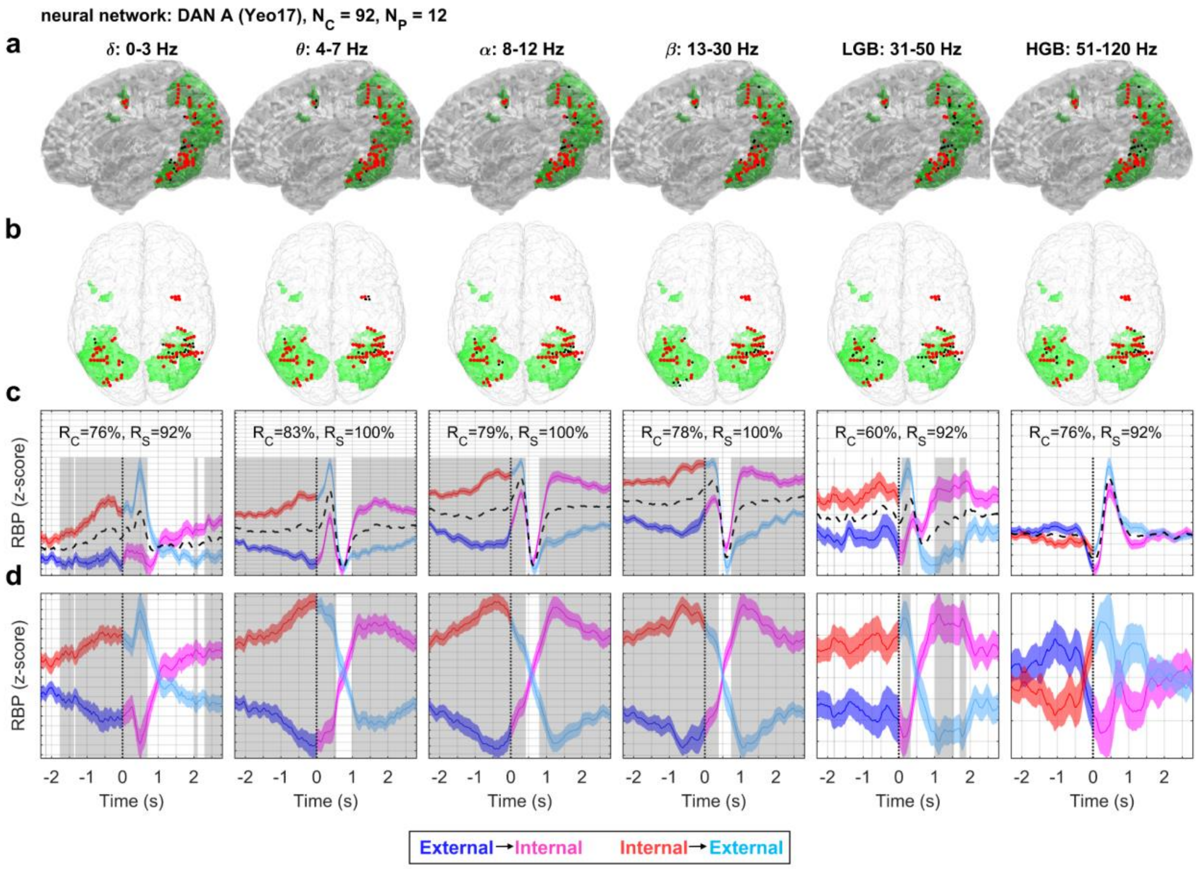


**Supplementary Figure S7. The DAN-A during the attention-switching task.** Same notations as in Fig. 4. The DAN-A (based on the Yeo-17 brain parcellation, highlighted in green) consisted of 92 channels in 12 subjects. The channels were anatomically distributed dominantly in the temporal lobe (fusiform gyrus, inferior and middle temporal gyrus), parietal lobe (angular gyrus and superior parietal lobule), occipital lobe (middle occipital gyrus) and a few channels in frontal lobe (middle frontal gyrus). The activations in LFB and LGB were consistent with overall DAN behavior. The HGB was dominated by a strong decrease (trough at t = 0 s), likely reflecting DAN-A deactivation during the button press (t = 0 s), followed by a rapid increase (peak at t = 500 ms) of power related to the presentation of the new visual stimulus for both tasks. On the network level, HGB was slightly more activated for the E-task than for the I-task (although the difference was not significant at the significance level $P$ = 0.001, FDR corrected).

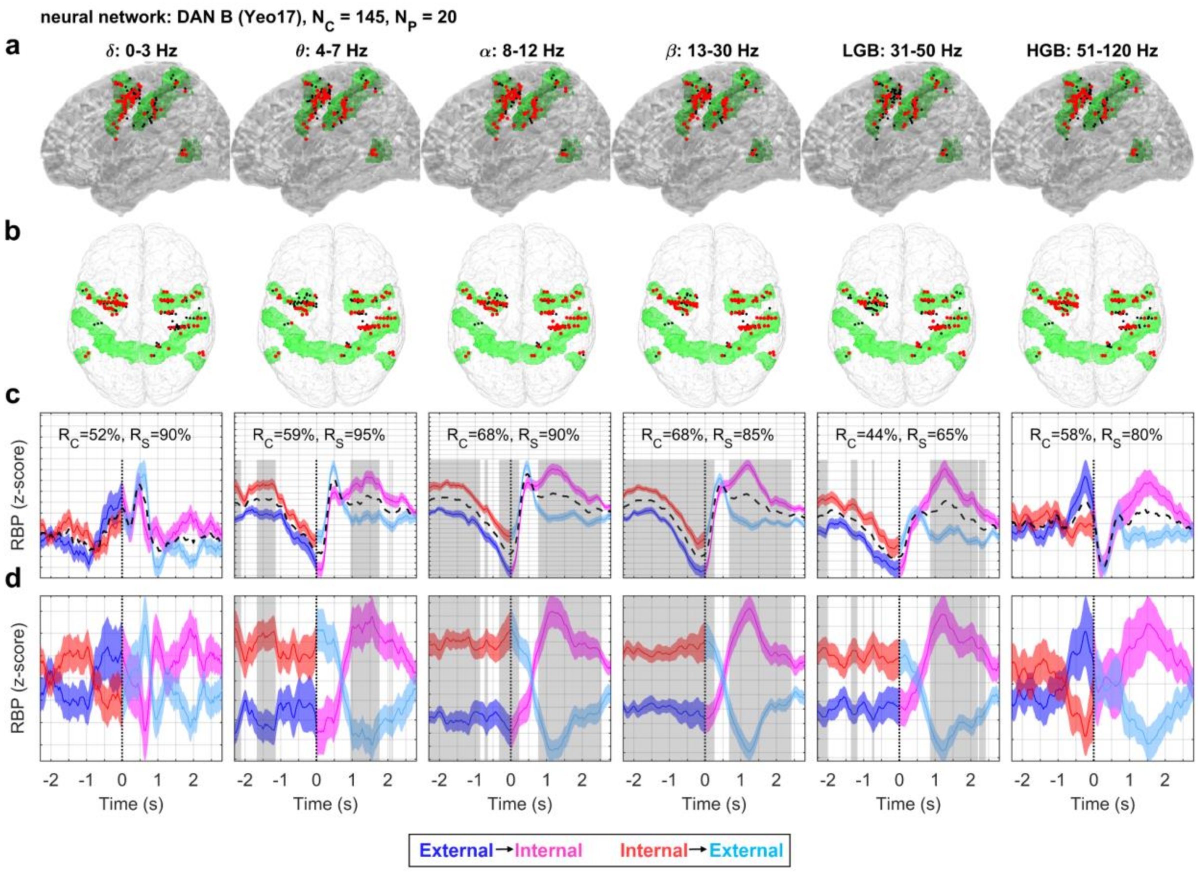


**Supplementary Figure S8. The DAN-B during the attention-switching task.** Same notations as in Fig. 4. The DAN-B (based on the Yeo-17 brain parcellation, highlighted in green) consisted of 145 channels in 20 subjects. The channels were anatomically distributed dominantly in the frontal lobe (middle and superior frontal gyrus, precentral gyrus), parietal lobe (postcentral gyrus, superior parietal lobule, and supramarginal gyrus) and a few channels in the temporal lobe (middle and inferior temporal gyrus). In the LFBs, there was a strong activity common to both task conditions (black dashed lines), presumably related to the button press and presentation of the new stimulus. After the subtraction of this common activity (bottom row), the polarity of the LFB activations were consistent with the overall behavior of the DAN. The HGB power showed an interesting modulation, with a peak in the E-task shortly before the stimulus presentation (blue curve; $t$ = –250 ms), a rapid decrease (trough at $t$ = 200 ms), followed by a gradual increase, which ceased for the I-task (cyan curve; $t$ = 700 ms), but continued for the E-task (magenta curve' peaking around $t$ = 1.5 s). However, the difference between the task conditions was not significant at the significance level $P$ = 0.001 (FDR corrected).

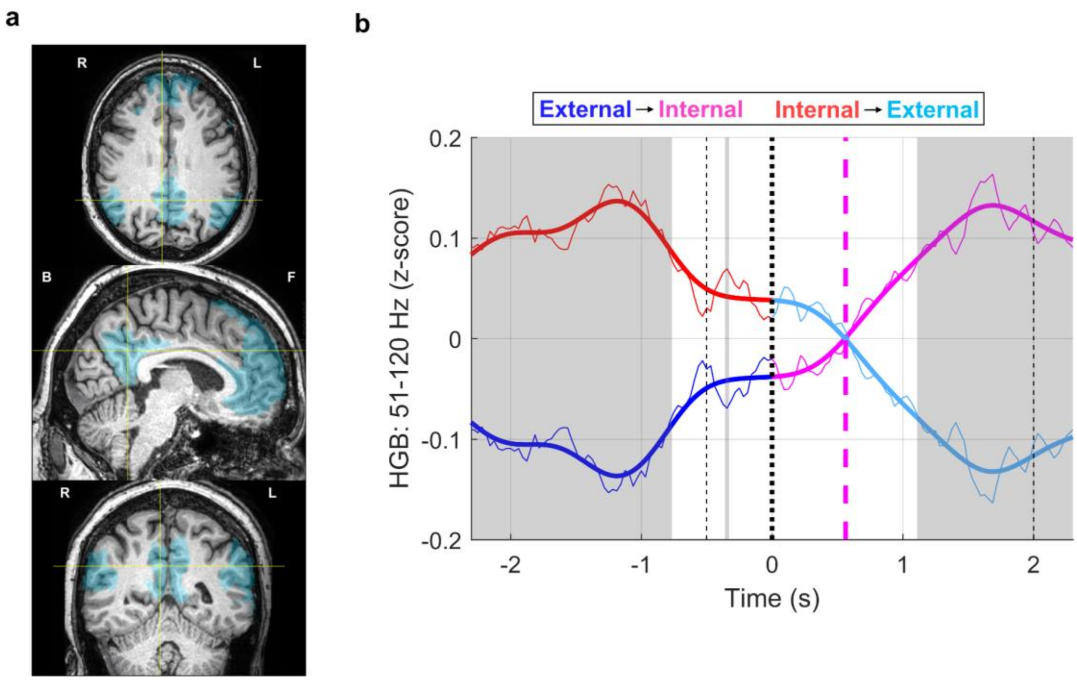


**Supplementary Figure S9. Crossover point of HGB activity during the attention-switching task. An** exemplary channel (subject P11) located in the posterior cingulate cortex, one of the major nodes of the DMN network, was selected for illustration. **(a)** Pre-implantation MRI (gray-scale) with the DMN highlighted in blue. Yellow crosshairs indicate the position of the selected channel. Top panel: axial view; middle panel: sagittal view; bottom panel: coronal view. R = right, L = left, F = front, B = back. **(b)** Corresponding, trial-averaged HGB power (51–120 Hz) after subtraction of the common (condition non-specific) signal for both task conditions: external–internal attention switching (blue–magenta) and internal–external attention switching (red–cyan). The thick curves are the HGB activity after low-pass (< 1 Hz) filtering, applied to smooth the jittered HGB activity. Gray background indicates significant activity ($P < 0.05$, FDR corrected), where the significance of difference was tested across different trials of the task conditions. Black dotted line indicates the time of the task switch ($t = 0$ s). Magenta dashed line indicates time of the crossover point ($t = 0$ s). We further demanded that the crossover point is located in the time interval $t \in [-0.5, 2]$ s (thin vertical dotted lines).

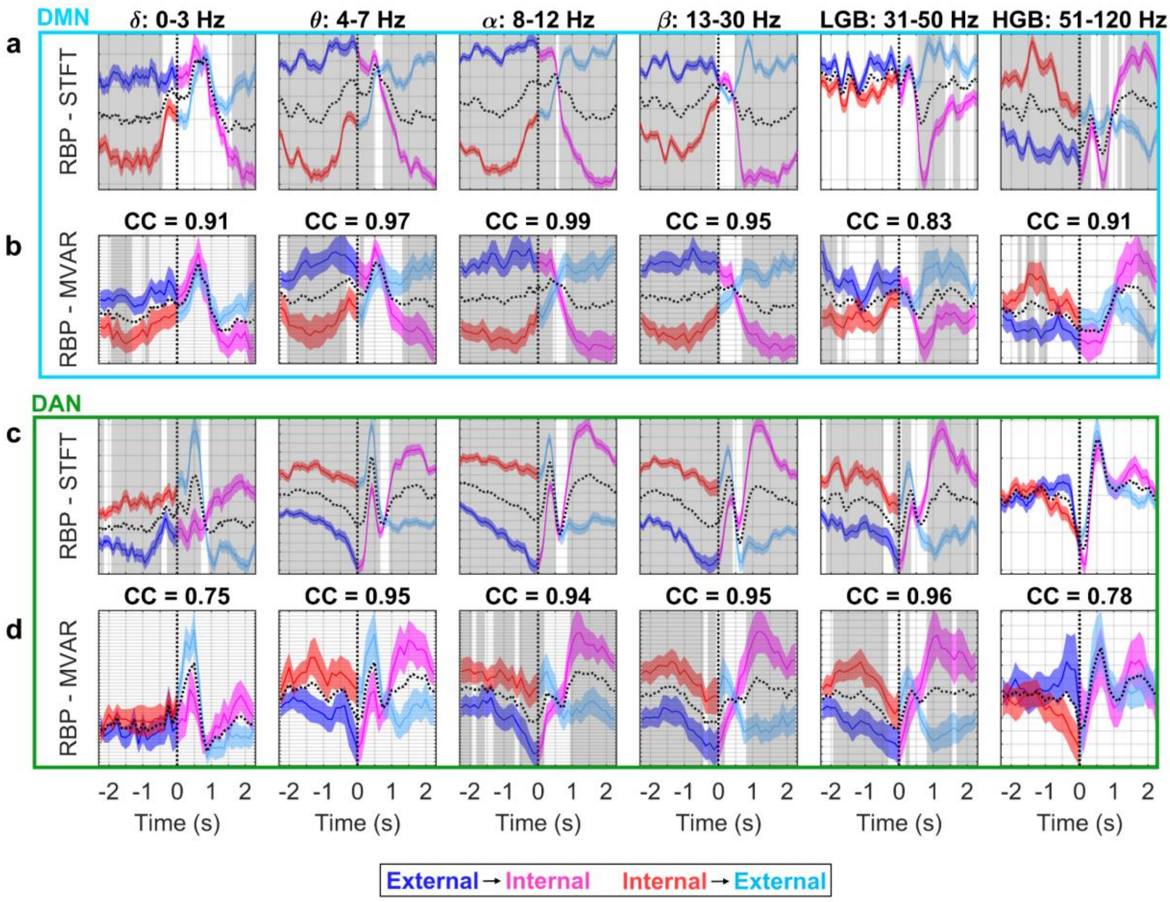


**Supplementary Figure S10. Comparison of RBP obtained by two different methods: STFT and an MVAR model.** The RBP is shown in the six different frequency bands (columns) indicated by the titles. The RBP was computed as a mean across the iEEG channels of the 17 subjects selected for the functional and effective connectivity presented in Fig. 7. Each subplot: same conventions as in Fig. 4. **(a)** RBP obtained by STFT for the DMN. **(b)** RBP obtained by the MVAR model for the DMN. The title above the second row in each subplot indicates the correlation coefficient (CC) between RBP of the two methods for both task conditions. **(c, d)** Same as in a and b but for the DAN. The overall CC was quite high (0.91 ± 0.02, mean ± SEM, across the six frequency bands and both networks), confirming that the MVAR model was able to capture the major power modulations observed with the more common STFT.
